# The perils of intralocus recombination for inferences of molecular convergence

**DOI:** 10.1101/393124

**Authors:** Fábio K. Mendes, Andrew Livera, Matthew W. Hahn

**Affiliations:** Department of Computer Science, The University of Auckland; Department of Biology, Indiana University; Department of Computer Science, Indiana University

## Abstract

Accurate inferences of convergence require that the appropriate tree topology be used. If there is a mismatch between the tree a trait has evolved along and the tree used for analysis, then false inferences of convergence (“hemiplasy”) can occur. To avoid problems of hemiplasy when there are high levels of gene tree discordance with the species tree, researchers have begun to construct tree topologies from individual loci. However, due to intralocus recombination even locus-specific trees may contain multiple topologies within them. This implies that the use of individual tree topologies discordant with the species tree can still lead to incorrect inferences about molecular convergence. Here we examine the frequency with which single exons and single protein-coding genes contain multiple underlying tree topologies, in primates and *Drosophila*, and quantify the effects of hemiplasy when using trees inferred from individual loci. In both clades we find that there are most often multiple diagnosable topologies within single exons and whole genes, with 91% of *Drosophila* protein-coding genes containing multiple topologies. Because of this underlying topological heterogeneity, even using trees inferred from individual protein-coding genes results in 25% and 38% of substitutions falsely labeled as convergent in primates and *Drosophila*, respectively. While constructing local trees can reduce the problem of hemiplasy, our results suggest that it will be difficult to completely avoid false inferences of convergence. We conclude by suggesting several ways forward in the analysis of convergent evolution, for both molecular and morphological characters.

## Introduction

Cases of phenotypic convergence represent some of the most striking and clear examples of the power of natural selection. Recent work has examined how often phenotypic convergence has been accompanied by convergent molecular changes in the same sites, genes, or pathways (reviewed in Rosenblum et al., 2014). While several good examples have been found (e.g., Christin et al., 2008; Manceau et al., 2010; Zhen et al., 2012; Natarajan et al., 2016), an equally striking pattern is the ubiquity of molecular convergence itself. Whole-genome comparisons have revealed high levels of molecular convergence in multiple clades (Bazykin et al., 2007; Rokas and Carroll, 2008; Foote et al., 2015; Goldstein et al., 2015; Zou and Zhang, 2015a; Chikina et al., 2016; Partha et al., 2017; Thomas et al., 2017). Although a large fraction of this convergence is undoubtedly due to non-adaptive processes (Thomas and Hahn, 2015; Zou and Zhang, 2015b) patterns of molecular convergence hold the promise of revealing general mechanisms of protein evolution (Storz, 2016).

A major issue with inferences of convergence is heterogeneity in the gene trees underlying the traits or characters of interest (Hahn and Nakhleh, 2016). Gene tree topologies (which are not necessarily in protein-coding genes) can differ from the species tree topology for multiple biological and technical reasons. The two major biological causes are introgression/hybridization and incomplete lineage sorting (ILS). Both introgression and ILS appear to be quite common across the tree of life, with massive amounts of gene tree discordance uncovered in both ancient and recent divergences (e.g., Pollard et al., 2006; The *Heliconius* Genome Consortium, 2012; Garrigan et al., 2012; Scally et al., 2012; Martin et al., 2013; Brawand et al., 2014; Jarvis et al., 2014; Fontaine et al., 2015; Novikova et al., 2016; Pease et al., 2016). Due to recombination along chromosomes, gene tree topologies can differ between neighboring nucleotides (Fig. 1a), with the similarity of topologies decaying on the scale of linkage disequilibrium if ILS is the only process acting (Slatkin and Pollack, 2006).

**Figure 1:**
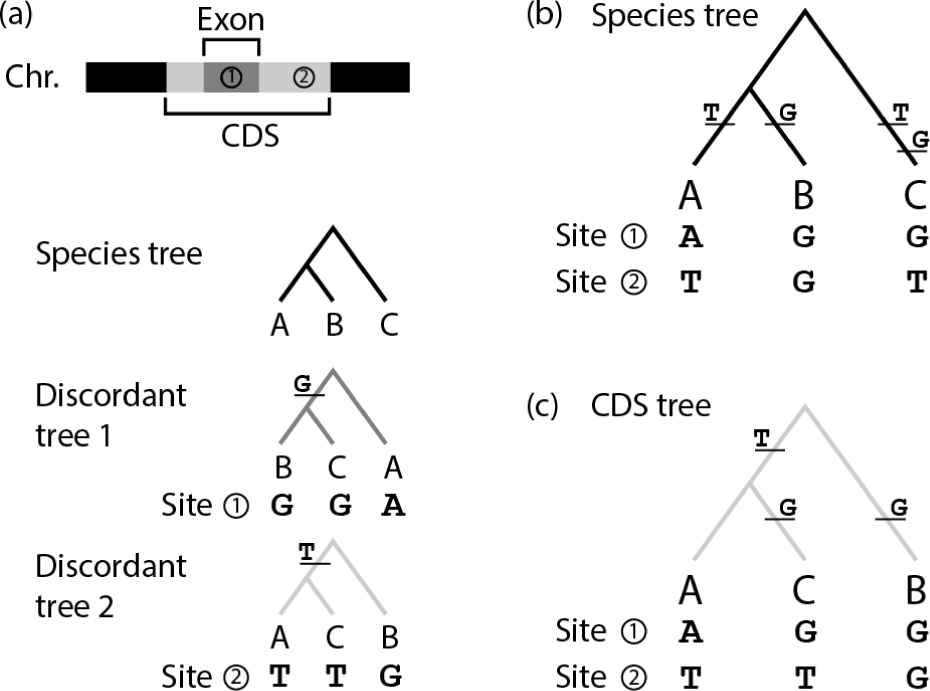
Hemiplasy on species trees and gene trees. Panel (a) shows substitutions that occur on each of the two discordant gene trees, and the corresponding site patterns produced. (b) Incorrectly inferred convergent substitutions (from the site patterns produced in [a]) when analyses are conducted on the species tree topology. (c) Incorrectly inferred convergent substitutions when analyses are conducted on the CDS tree (discordant tree topology 2).

Gene tree heterogeneity affects inferences of convergent evolution because single transitions along discordant topologies can be falsely inferred to involve multiple transitions when the data are analyzed using the species tree (Fig. 1b). This phenomenon has been dubbed “hemiplasy” (Avise and Robinson, 2008), in contrast to true biological homoplasy involving multiple transitions. Hemiplasy generates false signals of molecular convergence because substitutions along discordant internal branches – those that do not appear in the topology used for analysis – will appear to have occurred multiple times when these substitutions are mapped to the species tree (Fig. 1b). Molecular convergence due to hemiplasy does not reflect any biological property of the sites where it occurs, as it is solely the result of an artifact of using a tree for analysis that does not match the topology of the locus that generated the data. Hemiplasy has been explicitly identified in multiple datasets containing gene tree discordance (e.g., Mendes and Hahn, 2016; Mendes et al., 2016; Pease et al., 2016; Copetti et al., 2017; Palesch et al., 2018; Wu et al., 2018), and likely contributes to observed patterns of molecular evolution in many more datasets (e.g., Brawand et al., 2014; Neafsey et al., 2015; Thomas and Hahn, 2015; Zou and Zhang, 2015b).

In order to help control for hemiplasy, and to more accurately identify instances of homoplasy, several authors have constructed individual gene trees for every locus (Good et al., 2013; Copetti et al., 2017; Zou and Zhang, 2017). The aim of making gene trees is to use the same topology for analysis as the one that generated the data. This method will reduce the effect of hemiplasy if sites within a locus have evolved along its inferred gene tree topology, and indeed the use of individual gene tree topologies appears to have reduced the signal of false homoplasy relative to the case where the species tree is used for analysis (Copetti et al., 2017; Zou and Zhang, 2017). However, using individual gene trees will still not produce an accurate view of convergence if there is within-locus heterogeneity in the underlying topologies. As mentioned above, recombination can cause neighboring sites to have different topologies, such that the inferred topology from a region spanning multiple recombination breakpoints may include multiple underlying topologies, each incongruent with one another and possibly discordant with the species tree. In mammals it has been observed that different exons of the same protein-coding gene are no more likely to have similar topologies than exons from different genes, likely because of within-locus recombination (Scornavacca and Galtier, 2017). Therefore, while the use of individual gene tree topologies as the unit of analysis may reduce the problem of hemiplasy, it will not eliminate it. Instead, sites incongruent with the focal topology used for analysis will be falsely inferred to be homoplastic (Fig. 1c).

Here, we examine the degree of within-locus gene tree heterogeneity in two datasets, one from primates and one from *Drosophila*. Both of these datasets represent very recent splits among species, allowing us to minimize the effect of true homoplasy. We examine both within-exon and within-gene variation in tree topologies, quantifying the effect on hemiplasy of assuming that trees inferred from either partition represent the history of the whole locus. Due to intralocus recombination, results from both datasets reveal that false signals of convergence will be common even when locus-specific trees are inferred. We conclude by suggesting several ways that this phenomenon can be avoided.

## Methods

### *(a) Drosophila* data

Our analysis focuses on three closely related *Drosophila* species (*D. simulans*, *D. mauritiana*, and *D. sechellia*) and an outgroup (*D. melanogaster*). Aligned exonic sequences from 27 arthropod species were downloaded from the UCSC Genome Browser (Kent et al., 2002), and sequences of three of the species (*D. melanogaster*, *D. simulans*, and *D. sechellia*) were extracted from it. Exonic sequences were extracted from the *D. mauritiana* genome assembly using its gene annotation files (available from www.popoolation.at/mauritiana_genome/). Only the longest isoforms were kept. *D. melanogaster* exons were then used as a database against which those of *D. mauritiana* were compared (using BLAST, v.2.6.0+; Altschul et al., 1990) in order to establish orthology. Finally, the best matches of the BLAST search were added to the three-species alignments containing the corresponding *D. melanogaster* sequences, and all four sequences in each exon file were re-aligned using MUSCLE v.3.8.31 (Edgar, 2004) in its default setting. We removed exon alignments with more than 5% of their sites having missing data, and alignments to which a *D. mauritiana* sequence could not be assigned. A total of 35,050 exons remained after filtering. Protein-coding gene (“CDS”) alignments were then produced by concatenating exon alignments belonging to the same gene (totalling 10,353 CDS alignments).

### (b) Primate data

Our analysis focuses on three closely related primates (human, chimpanzee, and gorilla) and an outgroup (orangutan). Aligned exonic sequences from 20 primate species were downloaded from the UCSC Genome Browser, from which human, chimpanzee, gorilla, and orangutan sequences were extracted. We filtered out alignments with a human isoform other than the longest one, as well as alignments with more than 5% missing data in one or more species. The longest human isoform sequences were determined using the human refGene table (O’Leary et al., 2015) obtained from the UCSC Table Browser (Karolchik et al., 2004). A total of 147,428 exons remained after filtering. Again, exon alignments belonging to the same genes were concatenated, totalling 16,090 CDS alignments.

### (c) Site pattern tabulation and gene tree estimation

Each site in an alignment exhibits a site pattern, which consists of a specific series of nucleotides (one nucleotide per species) appearing in some fixed order. In our datasets, site patterns were considered informative if exactly two ingroup species shared a derived state (with the ancestral state being the nucleotide present in the outgroup). Site patterns were tabulated for all exon and CDS alignments, according to their informativeness and congruence status. Site patterns were classified as “concordant” if they could be produced by a single substitution on a species tree internal branch (via maximum-parsimony mapping), and as “discordant” otherwise (Mendes and Hahn, 2018). Site patterns are defined here as “congruent” if they could be produced by a single substitution on a gene tree internal branch (regardless of whether the gene tree is concordant with the species tree), and as “incongruent” otherwise. Gene trees were estimated for all exon and CDS alignments using IQ-TREE v.1.6.6 under the corresponding best nucleotide model (Nguyen et al., 2015), and rooted with D. melanogaster and orangutan in the *Drosophila* and primate data sets, respectively. Only trees estimated from alignments with at least one informative site pattern were considered in these analyses.

## Results

### (a) Informative sites across protein-coding genes in primates and Drosophila

Both of the datasets considered here have three ingroup taxa, and therefore three possible rooted topologies and three informative site patterns (two of which are shown in figure Fig. 1a). In primates, the accepted species tree groups chimpanzees and humans together, with gorillas sister to these two species (Scally et al., 2012); we will denote this topology as simply “(ch,hum)”. The two discordant topologies group either chimpanzees and gorillas (ch,gor) or humans and gorillas (hum,gor). The species tree topology among the *Drosophila* species studied here is less clear, though the preponderance of evidence supports the grouping of *D. simulans* and *D. sechellia* first (sim,sec), with *D. mauritiana* sister to both of these (Garrigan et al., 2012; Pease and Hahn, 2013). A maximum-likelihood topology inferred from concatenating all of the data used here also found this topology. The two discordant topologies are (sim,mau) and (sec,mau).

We looked within protein-coding genes for variable sites (both synonymous and nonsynonymous) that supported these different topologies. In total, we used 24,468,005 bp of aligned sequence in primates and 13,497,746 bp of aligned sequence in *Drosophila*. Given the recent divergences between species in both clades (less than 2% divergence between all pairs of ingroup species; Garrigan et al., 2012; Scally et al., 2012), we expect that the vast majority of informative sites represent a single substitution rather than multiple substitutions. As a consequence, we can use informative site patterns to tell us about the underlying gene tree topologies and their relative frequencies.

The number of informative site patterns varied among exons and among whole protein-coding sequences (CDSs). In both categories the most common observation was a total lack of informative sites: this was true for both Drosophila and primates (Fig. 2). The most extreme case occurs within exons among primates, where approximately 80% of all single exons do not contain a single informative substitution (Fig. 2a). Due to larger effective population sizes and higher overall levels of (ancestral) polymorphism – as well as longer exon lengths (see next section) – only 43% of *Drosophila* single exons completely lack any informative sites (Fig. 2a). Nevertheless, results from both species demonstrate why researchers often need to combine exons together when estimating tree topologies. Indeed, the fraction of whole coding sequences that lack any informative sites drops to approximately 25% and 12% for primates and *Drosophila*, respectively (Fig. 2b). At the other extreme, single exons regularly contain 5 or more informative sites (Fig. 2a), and single CDSs can contain upwards of 50 informative sites (Fig. 2b). Any exon or CDS containing multiple informative sites can possibly contain diagnosably different tree topologies. Therefore, we next looked at the genome-wide frequencies of the three site patterns supporting different topologies.

**Figure 2:**
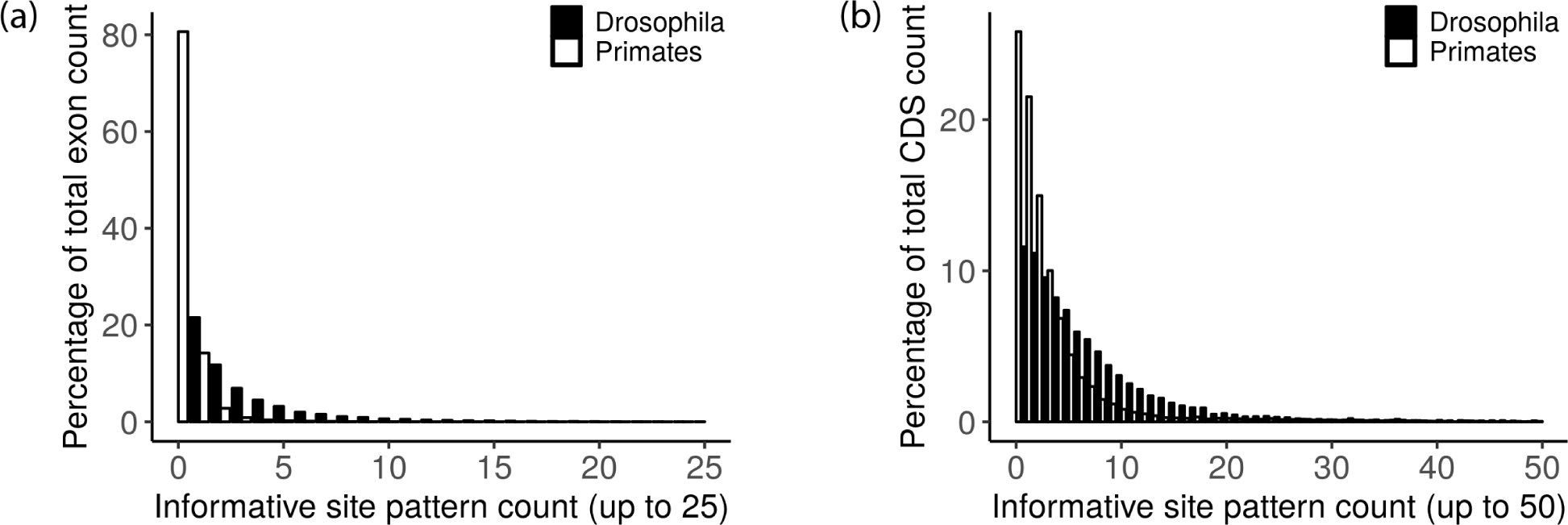
Percentage of single (a) exons or (b) CDSs that have a given number of informative sites.

**Figure 3:**
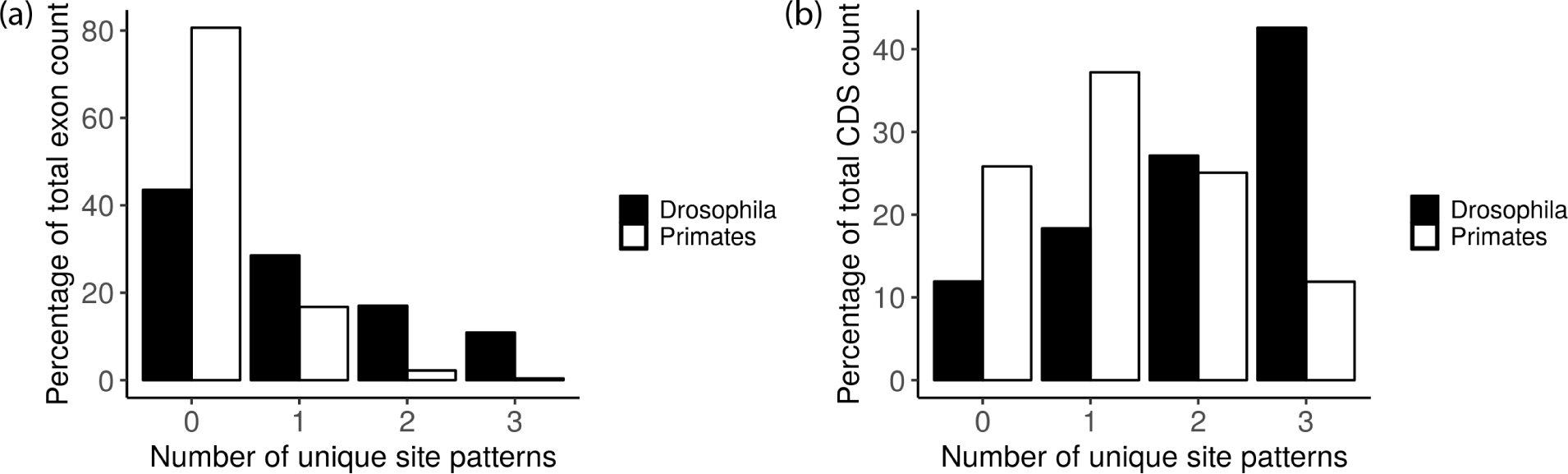
Percentage of (a) exons or (b) CDSs containing a given number of unique site patterns within them.

We found high levels of gene tree discordance in both clades. As expected in primates, the most common site pattern grouped chimpanzees and humans: this pattern was present among 46.9% of informative sites (out of 73,005 informative sites in total). In *Drosophila*, the pattern grouping D. simulans and D. sechellia was present in 37.0% of sites (out of 65,269 informative sites in total), again suggesting that this topology is the most common tree and likely represents the species tree. It is also evident that in both datasets site patterns discordant with the species tree account for more than 50% of all informative substitutions. This initial result implies that, if the species tree were used for analysis, more than half of all substitutions shared between two non-sister species would be falsely inferred to be due to homoplasy.

### (b) Gene tree heterogeneity within exons and whole coding sequences

In both primates and *Drosophila*, the results thus far demonstrate that there is a large amount of underlying gene tree discordance and the potential for multiple gene trees to exist (and be diagnosed) for single exons and CDSs. If recombination occurs within exons, then we should be able to observe multiple incongruent site patterns within a single exon. Likewise, recombination either within or between exons (i.e., in introns) will mean that different site patterns can be observed within a single CDS, indicating the presence of multiple underlying gene trees.

We observed evidence for gene tree heterogeneity within both exons and whole coding sequences (CDSs), in both primates and *Drosophila*. The average length of all exons examined here is approximately 166 bp in primates and 385 bp in *Drosophila*. Nevertheless, we still found two or more unique informative site patterns in 2.5% of primate exons and 28% of *Drosophila* exons (Fig. 3a). As a large fraction of exons have no informative site patterns in both clades, it may be more useful to express these numbers as a percentage of all exons containing two or more informative sites, such that gene tree heterogeneity can be assessed. We observe two or more incongruent site patterns in 50% of all primate exons and 80% of all *Drosophila* exons containing at least two informative site patterns.

The effects of recombination are more pronounced for whole coding sequences, as there is more opportunity for recombination (especially within introns) to bring together two or more topologies. Indeed, approximately 36% of primate CDSs and 70% of *Drosophila* CDSs have two or more unique site patterns within them (Fig. 3b). Expressed as a proportion of CDSs with at least two informative sites (regardless of whether they are incongruent with each other), these numbers jump to 70% and 91% of CDSs for primates and *Drosophila*, respectively. In both cases, the most common observation across the genome will be for single protein-coding genes to contain either two or three conflicting tree topologies within them. This scenario is even more common than for them to contain only a single diagnosable topology (Fig. 3b).

These results demonstrate that even very small genomic regions – such as exons – may contain multiple gene trees within them. This implies that we may misidentify convergent substitutions when inferring local trees. In the next section we attempt to quantify the extent to which this can occur, and how the effects can differ among topologies.

### (c) Consequences for inferences of convergence

In order to avoid the problems associated with hemiplasy, we would like to use the same tree that the data evolved along for our analysis. When there is a mismatch between these trees, we may still be falsely inferring more convergent evolution than is actually occurring. If we assume that in our dataset all informative substitutions reflect their underlying gene tree topologies (see Discussion for caveats associated with this assumption), then we can use the topologies inferred from different genomic compartments to quantify the effects of hemiplasy within each.

As mentioned earlier, in neither primates nor *Drosophila* does the site pattern matching the presumed species tree make up a majority of informative sites. In *Drosophila* only 37.0% of sites match the species tree (sec,sim), with 33.1% joining *D. sechellia* and *D. mauritiana* (sec,mau) and 29.9% joining simulans and *D. mauritiana* ([sim,mau]; Fig. 4a). As a consequence, if the species tree were used for analysis we would falsely infer 63% of informative sites to have evolved convergently. Similarly, in primates only 46.9% of informative sites match the species tree (ch,hum), with 32.7% joining chimpanzee and gorilla (ch,gor) and 20.4% joining gorilla and human ([gor,hum]; Fig. 4b). In this case approximately 53% of all informative sites would be falsely inferred to have evolved convergently if the species tree were the sole tree used for analysis (i.e., these are hemiplastic rather than homoplastic).

Instead of using the species tree to analyze all the data, we can instead infer trees for each exon or protein-coding gene. To a first approximation, the trees inferred in these cases represent the most common informative site pattern at each locus, while still hiding the presence of incongruent site patterns within them. Using longer loci to infer trees means that proportionally more individual topologies can be inferred (because there are more informative sites per locus; Fig. 2), but their use also results in the combination of more incongruent sites within a locus (Fig. 3). In total, we were able to infer 28,529 (primates) and 19,783 (*Drosophila*) trees from exons and 11,934 (primates) and 9,119 (*Drosophila*) trees from CDSs. This represents 19.3% and 56.4% of all exons and 74.1% and 88% of all CDSs in the dataset (after filtering, see Methods) in primates and *Drosophila*, respectively.

**Figure 4:**
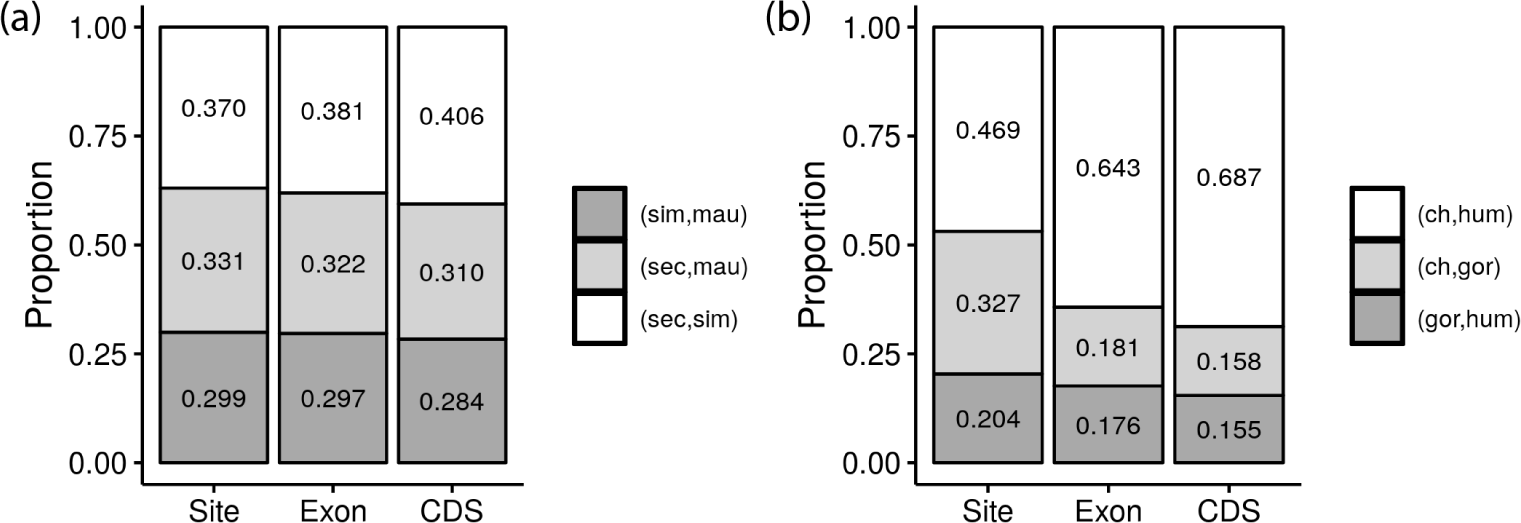
The proportion of inferred trees supporting each of the three possible topologies in (a) *Drosophila* and (b) Primates. Each site in the "Site" category is assigned a tree according to its maximum-parsimony mapping.

As expected, in both datasets the most common site-type matches the most common inferred tree, from both exons and CDSs (Fig. 4). Also as expected, larger bins (i.e., CDSs) contain more informative sites and are therefore each more likely to individually contain more sites supporting the species tree, resulting in a higher fraction of inferred trees with this topology (Fig. 4). Although we were able to infer proportionally fewer topologies from individual exons (containing fewer aligned positions in total), in both primates and *Drosophila* the percentage of inferred trees from exons supporting the species topology more closely approaches the true number (i.e., the number implied by the site counts; Fig. 4).

Using the inferred trees from CDSs and exons, we can begin to ask how often hemiplasy would occur within each compartment if analyses were conducted using these trees. Based on the results above, we expect that hemiplasy will be a larger problem for CDSs, as they bring together more unique topologies within a single locus. In order to quantify the effect of hemiplasy, for each inferred tree we count the number and proportion of all informative sites within them that are incongruent. Regardless of whether an inferred tree is concordant or discordant with the species tree, sites incongruent with it would result in hemiplastic substitutions (Fig. 1c).

Using whole protein-coding genes (CDSs) as the unit of analysis, we find that inferred gene trees contain on average 24.6% and 38.3% incongruent sites for primates and *Drosophila*, respectively. In other words, in a single gene inferred to have a particular topology in primates, 24.6% of the informative sites within it would not support this topology. While this is an improvement compared to using the species tree for all loci (where 53% and 63% of all sites were incongruent for primates and *Drosophila*), this still implies a high level of hemiplasy. There is some variation among topologies, with primate gene trees matching the species tree containing 21.7% incongruent sites, and the two discordant trees contain 33.0% (gor,hum) and 29.1% (ch,gor) incongruent sites (Fig. 5a). Similar patterns among topologies are observed in *Drosophila* (Fig. 5c).

**Figure 5:**
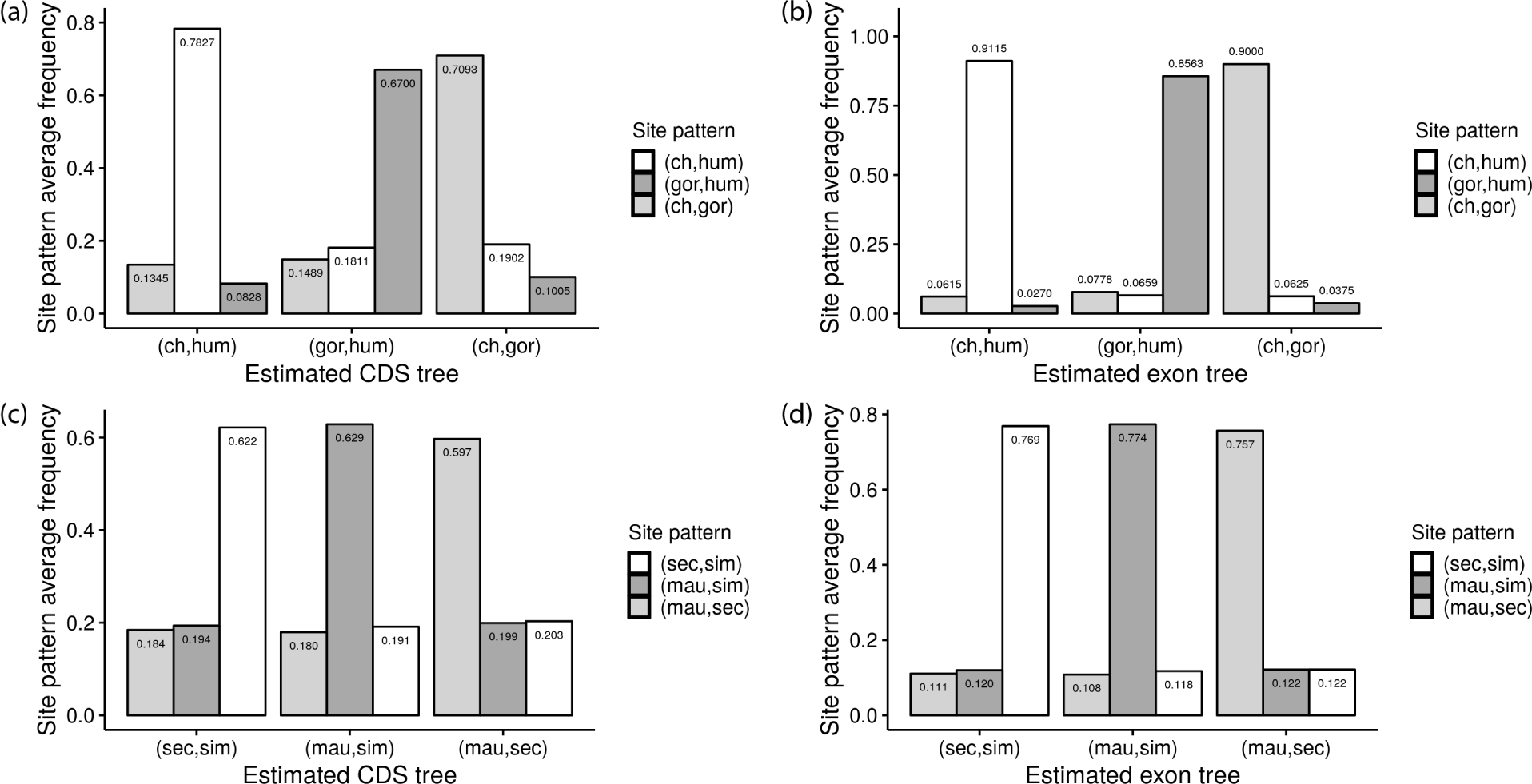
Average frequency of site patterns within inferred tree topologies. For each CDS or exon in which we could infer a tree topology, the frequency of informative site patterns of each type was calculated. Site patterns were classified according to the tree they support using a maximum-parsimony mapping scheme. (a) Primate CDSs (n=11,934), (b) Primate exons (n=28,529), (c) *Drosophila* CDSs (n=9,119), and (d) *Drosophila* exons (n=19,783). Note that the site pattern supporting (sec,sim) is slightly more common than (mau,sim) in the rightmost comparison in panel d.

If instead we use exons to infer topologies, the resulting gene trees contain on average 10% and 23.3% incongruent sites for primates and *Drosophila*, respectively. This is again an improvement compared to the use of CDSs, though there would still be a large number of hemiplastic substitutions (and a much smaller number of total sites available for analysis). There is also again some variation in the number of incongruent sites among topologies, with between 8-14% and 22-24% incongruent sites for primates and *Drosophila*, respectively (Fig. 5b and 5d).

Among the tree topologies inferred from individual loci, we do not expect the frequency of the two incongruent sites to necessarily be the same within each. For discordant trees the frequencies of the two incongruent site patterns contained within them are expected to be unequal, as incongruent concordant sites will be (almost by definition) more common than incongruent discordant sites. This pattern is generally observed, as more incongruent sites within discordant gene trees supported the concordant topology in seven of the eight possible comparisons (i.e., two discordant trees, in CDSs and exons, for primates and *Drosophila*; Fig. 5). Individual discordant gene trees will therefore result in the over-representation of a specific pattern of hemiplasy: the apparent convergence of states between the two taxa that are sister to each other in the species tree.

## Discussion

Convergence – or more generally, homoplasy – is inferred when multiple character-state transitions occur on a tree. However, convergence can be incorrectly identified when the tree a trait has evolved along is not the tree used for analysis (Avise and Robinson, 2008; Hahn and Nakhleh, 2016). To avoid such problems of “hemiplasy,” researchers are beginning to shift from using the species tree for all analyses to using individual gene trees inferred from each locus (e.g., Mendes and Hahn, 2016; Copetti et al., 2017; Ogilvie et al., 2017; Zou and Zhang, 2017). The hope is that gene trees inferred from smaller segments of the genome are more reflective of the local topology, such that nucleotide or amino acid substitutions in these segments are more accurately modeled by local trees. However, due to recombination, even trees inferred from very short stretches of the genome can include multiple underlying topologies. Here we have quantified the extent to which the construction of local topologies can mitigate the problems associated with hemiplasy, for two different datasets.

Our results suggest that constructing local trees can greatly reduce the problem of hemiplasy, though using trees from whole protein-coding sequences was much less effective than using trees from individual exons. Eukaryotic protein-coding sequences, which often include multiple exons and long introns, can contain multiple different tree topologies (Fig. 3). In fact, different exons in the same protein-coding gene may not share any more similarity in their topologies than do exons in different genes (e.g., Scornavacca and Galtier, 2017). Because of this underlying heterogeneity, inferred gene trees may contain incongruent sites due to hemiplasy and not homoplasy. Note that this inference requires that we have a more general definition of hemiplasy, which has often been applied only in relation to the species tree. Hemiplasy as defined here is the production of a homoplasy-like event when traits that have evolved along one topology are analyzed using a different topology (whether or not this is the species tree). We believe that this definition will help to clarify cases in which hemiplastic substitutions may still contribute to observed patterns of homoplasy, even when individual gene trees have been inferred (cf. Copetti et al., 2017; Zou and Zhang, 2017).

We observed relatively large differences in the proportion of incongruent sites within both exons and CDSs between primates and *Drosophila*, such that there were more than double the number of such sites in *Drosophila* exons relative to primate exons (Fig. 5). The biological reasons for such differences are manifold, likely including differences in mutation rates, recombination rates, and effective population sizes. But there are proportionally more incongruent sites genome-wide in the *Drosophila* dataset used here, which means that the benefit of inferring local trees should not be measured by the absolute number of incongruent sites. In fact, the construction of trees from smaller segments appears to have approximately equal efficacy in the two species: there are 53% and 63% incongruent sites for primates and *Drosophila* genome-wide, which reduces to 25% and 38% for inferred CDS trees, which further reduces to 10% and 23% for exon trees. This relatively consistent reduction in the number of sites affected by hemiplasy in the two datasets suggests that the construction of local trees will also work in other datasets.

Our estimates of the absolute number of sites affected by hemiplasy rest on the assumption that informative site patterns reflect only underlying tree topologies, and that there is no effect of true homoplasy. While both datasets contain very closely related species – with very low levels of nucleotide divergence – there is still a possibility that homoplasy has occurred. In order to estimate the proportion of incongruent sites due to homoplasy in any dataset, we have developed a population genetic model of both hemiplasy and homoplasy (Guerrero and Hahn, 2018). Using estimates of the speciation times, mutation rate, and effective population sizes of the primates studied here, this model predicts that 84% of sites incongruent with the species tree are due to hemiplasy, with the other 16% due to homoplasy. While there are a number of caveats to this estimate, if we take it as approximately correct it implies that the true proportion of hemiplastic sites in primates is 21% for CDSs (=25%*0.84) and 8.4% for exons (=10%*0.84). Unfortunately, we do not have such a prediction for the *Drosophila* data. While an estimate of this fraction may help to further correct numbers from *Drosophila*, the exact value will not affect the general conclusion that aggregating together segments of the genome with different histories results in incorrect inferences about homoplasy.

The results presented here may be of relevance to an ongoing debate surrounding the application of gene tree-based methods to the inference of species trees (Gatesy and Springer, 2014; Roch and Warnow, 2015; Edwards et al., 2016; Springer and Gatesy, 2016). It has been shown that concatenating together loci with different underlying trees can result in the inference of an incorrect species tree when using maximum likelihood methods (Kubatko and Degnan, 2007), though interestingly not when using distance-based methods or parsimony (at least for four-taxon rooted trees; (Liu and Edwards, 2009; Mendes and Hahn, 2018). One solution to this problem is to use methods that combine information from independently inferred gene trees to construct the species tree (e.g., Kubatko et al., 2009; Liu et al., 2010; Mirarab and Warnow, 2015). An issue with such methods is that they are only guaranteed to be more accurate than concatenation when each gene tree represents the topology from a non-recombining locus (Roch and Warnow, 2015). If gene trees are inferred from recombining loci, the assumptions of gene tree methods are violated (Gatesy and Springer, 2014; Springer and Gatesy, 2016). Our results support the contention that whole protein-coding loci (and individual exons) often include multiple different underlying topologies, and that this leads to biases in the proportion of gene trees supporting different topologies (Fig. 4). While these biases are generally quite small, we have only considered incomplete lineage sorting among three taxa – it is possible that with more taxa the problem becomes worse. Simulations conducted with many more sampled lineages revealed few errors when concatenating loci (Lanier and Knowles, 2012) though this and similar results (e.g., Bayzid and Warnow, 2013; Zimmermann et al., 2014) cast doubt on whether we should be worried about using concatenation for inferring the species tree at all.

Regardless of the methods used for species tree inference, it is clear that gene tree heterogeneity results in hemiplasy. The ideal scenario for avoiding problems of hemiplasy would be to infer trees from single sites, but this is infeasible when there are deeper splits in the tree because in these cases site patterns do not necessarily reflect underlying topologies. Another possible solution would be to only use trees from longer segments with little recombination, such as mtDNA or loci from low-recombination regions. However, this approach would both strongly limit the number of genes that could be studied and may not completely obviate the problems of false convergence. This is because incongruence between trees may be due to either biological or technical reasons. Here we have only addressed biological issues, but using an incorrect tree because of errors in inference will still result in hemiplasy (e.g., Goldstein et al., 2015; Mendes et al., 2016). The possibility of getting highly supported, yet incongruent, trees from a single mtDNA genome (e.g. Richards et al., 2018) suggests that technical issues may have large effects on hemiplasy in many contexts. A more promising approach may be to use recombining loci in a model-based framework that explicitly includes recombination. For instance, Bayesian methods like the one implemented in StarBEAST2 (Ogilvie et al., 2017) account for among-locus topological heterogeneity when inferring substitution rates by modeling gene tree discordance. Such methods employ the multispecies coalescent model, and could be applied to recombining loci if information about recombination rates is available; under the multispecies coalescent, the likelihood of a discordant gene tree and a single substitution (hemiplasy) can be compared to that of a concordant tree and multiple substitutions (homoplasy) without user input. With an extra step of stochastic mapping (Nielsen, 2012; Vaughan et al., 2014), the posterior distribution of all substitutions can be obtained. If recombination rates are unknown, Bayesian model comparison is one potential way to determine the optimal locus length for minimizing hemiplasy. In cases like these there will be a trade-off between the amount of signal per locus (i.e., how well each estimated gene tree will reflect the true gene tree) and how many different topologies are included within single loci.

Unfortunately, such model-based solutions are not currently available for researchers to use. What, then, is a way forward with today’s datasets? One partial solution we have previously proposed (Mendes et al., 2016) is to distinguish between different types of convergence, as some patterns cannot be caused by gene tree heterogeneity. Briefly, only convergent evolution from the same ancestral state in multiple lineages can be mimicked by hemiplasy; convergence arising from different ancestral states is not prone to hemiplasy (see Fig. 2 in Mendes et al., 2016). Therefore, even if an incongruent topology is used for analysis, inferences of convergent changes from different ancestral states will be immune to the problems we have discussed here. While such methods may limit the number of nucleotide sites available for analysis, they also greatly extend the types of characters that can be studied: they can be used even for traits where the gene tree is not known. Although the focus in this paper has been on molecular convergence, hemiplasy can affect any character, whether molecular or morphological, discrete or continuous (Hahn and Nakhleh, 2016; Mendes et al., 2018). For most such characters it will not be possible to infer the underlying tree topology or topologies, making methods that do not rely on knowing the gene tree much more generally applicable.

## Authors’ contributions

F.K.M and M.W.H. conceived the study, F.K.M. and A.L conducted analyses, F.K.M. and M.W.H. wrote the manuscript.

